# Identification of CD73 as the antigen of an antigen-unknown monoclonal antibody established by exosome immunization, and its antibody-drug conjugate exerts an antitumor effect on glioblastoma cell lines

**DOI:** 10.1101/2021.09.30.462527

**Authors:** Takahiro anzai, Shinji Saijou, Hiroki Takashima, Misato Hara, Shingo Hanaoka, Yasuhiro Matsumura, Masahiro Yasunaga

## Abstract

Development of antibodies against the native structure of membrane proteins with multiple transmembrane domains is challenging because it is difficult to prepare antigens with native structures. Previously, we successfully developed a monoclonal antibody against multi-pass membrane protein TMEM180 by exosome immunization in rats. This approach yielded antibod-ies that recognized cancer-specific antigens on the exosome. In this study, we performed im-munoprecipitation using magnetic beads to identify the antigen of one of the rat antibody clones, 0614, as CD73. We then converted antibody 0614 to human chimeric antibody 0614-5 and found that the humanized antibody did not inhibit CD73 enzymatic activity. Glioblastoma (GB) was the cancer type with the highest expression of CD73 in the tumor relative to healthy tissue. An antibody–drug conjugate (ADC) of 0614-5 exerted an antitumor effect on GB cell lines according to expression of CD73. The 0614-5-ADC could be used to treat cancers with high CD73 expression. In addition, our strategy could be used to determine the antigen of any antibody produced by exosome immunization.

## 1. Introduction

There is considerable motivation to identify new target molecules for the diagnosis and development of new cancer drugs. In previous work, we identified a new colorectal cancer tumor marker multi-pass membrane protein, TMEM180, which is also secreted in exosomes [1]. Development of antibodies against the native structure of membrane proteins with multiple transmembrane domains is challenging because it is difficult to pre-pare antigens with native structures. To overcome this difficulty, we successfully developed a TMEM180 monoclonal antibody by immunizing exosomes, which are TMEM180-positive [1]. On the other hand, because of exosome immunization, although the antigen is not TMEM180, antibody clones that recognize unknown antigens with high cancer specificity have also been established. By determining the antigen of an antibody, it is possible to obtain an antibody against a membrane protein. In this study, we identified the antigen of a cancer-specific recognition monoclonal antibody as CD73 and characterized the antibody. We identified glioblastoma (GB) as the cancer with the highest cancer-specific expression of CD73 and demonstrated that an antibody–drug conjugate of the anti-CD73 monoclonal antibody exerted an antitumor effect on our GB cell lines.

## 2. Results and discussions

### 2.1. The target antigen of antibody clone 0614 is CD73

We developed antibodies by exosome immunization, as described previously [1]. Briefly, we immunized rats with tumor-derived exosomes purified from the supernatant of the colon cancer cell line DLD-1. We then analyzed supernatants of hybridoma cells containing antibodies by flow cytometry and screened both DLD-1 colorectal cancer (CRC) cells (positive) and K562 myeloma cells (negative) (Fig. S1). We established a rat IgG2a clone, 0614, as an antibody of unknown antigens. To identify the antigen, we isolated and purified it from exosomes (Fig. 1A). In this approach, the target antibody is first biotinylated and bound to avidin-conjugated magnetic beads. Then, these beads are bound to an antigen-positive exosome. An exosome–antibody complex is isolated by immunoprecipitation. After the antigen is dissociated from magnetic beads, it is detected by SDS-PAGE and western blotting (Fig. 1A). We then performed immunoprecipitation of exosomes derived from DLD-1 cells using biotinylated antibody 0614 and avidin-bound magnetic beads, and analyzed the resultant samples by SDS-PAGE (Fig. 1B) and western blotting (Fig. 1C). We succeeded in obtaining an antigen candidate of about 70 kDa. To al-low more accurate antigen determination, the samples were separated by two-dimensional electrophoresis (Fig. 1D). Mass spectrometry revealed 5’-nucleotidase (UniProt:P21589) and heat shock 70 kDa protein 1A (UniProt:P0DMV8) and 1B (Uni-Prot:P0DMV9) as candidates. Because the antigen of 0614 can be detected by flow cytometry, we focused on 5’-nucleotidase, also called CD73. We constructed and purified the extracellular domain of CD73 [2–4] with a C-terminal His-tag as a recombinant protein (Fig. 1E), and then performed western blotting; this analysis confirmed that the antigen of antibody 0614 is CD73 (Fig. 1F). In functional terms, CD73 is an enzyme that converts phosphorylated adenosine AMP; it is localized on the membrane surface of regulatory T cells and cancer cells and activates an immune response against adenosine [5,6]. There are some reports that CD73 expressed on exosomes has enzymatic activity, and exosome-derived adenosine results in T cell inhibition and impairs the antitumor immune response [7–9]. Thus, it is reasonable that anti-CD73 antibody could be established by exosome immunization, we decide to proceed with development of 0614 antibody that recognizes CD73 on exosomes.

**Figure 1.**
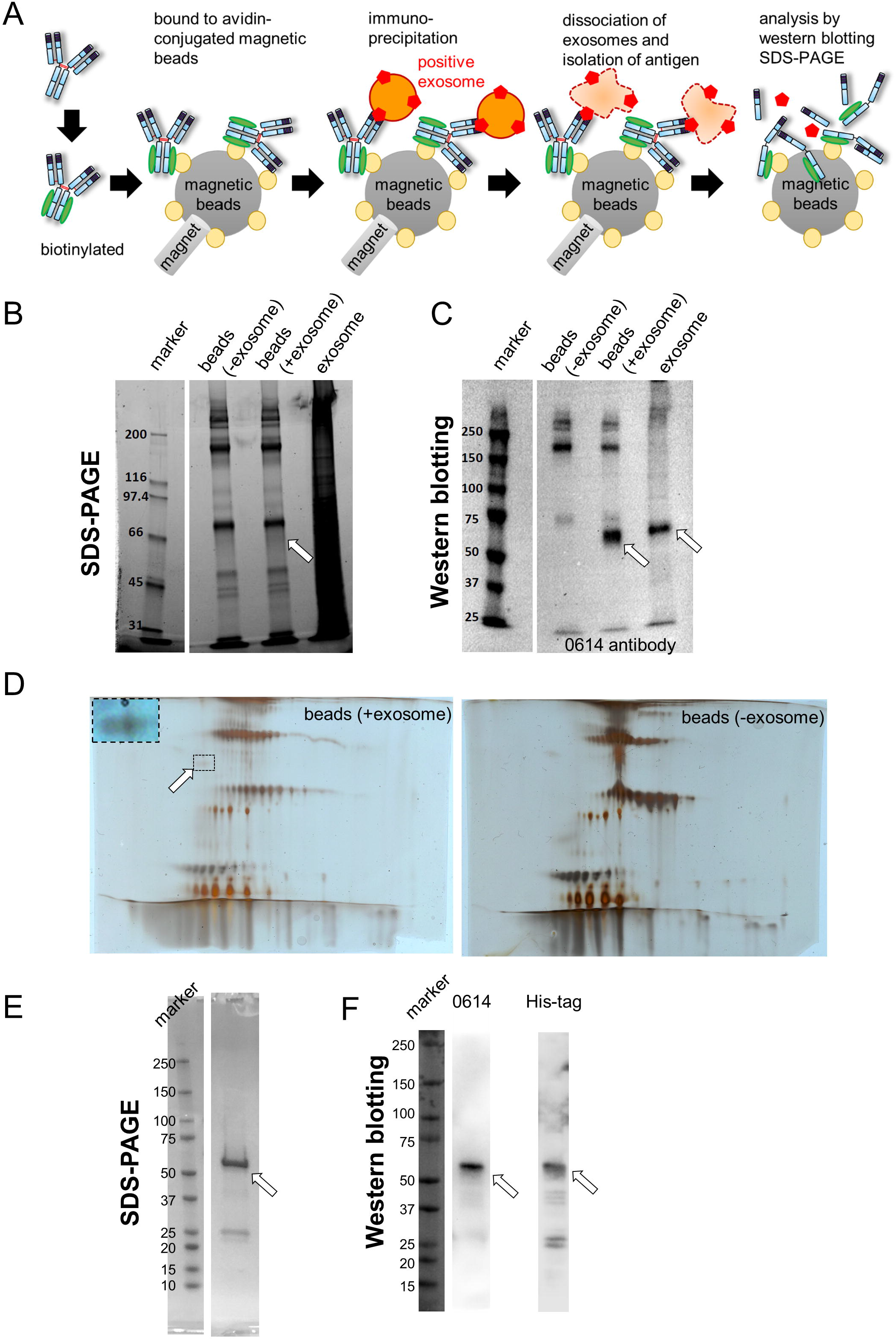
Identification of CD73 as the antigen of antibody 0614. A. Schema for identification of the unknown antigen of antibody 0614. B, C. SDS-PAGE (B) and western blot (C) analyses of samples immunoprecipitated with avidin magnetic beads conjugated to biotinylated antibody 0614. Target antigen containing exosome samples was loaded as a positive control. The position of the target antigen is indicated by a white arrow. D. 2D-PAGE analysis of samples immunoprecipitated with avidin magnetic beads conjugated to biotinylated antibody 0614. The position of the target antigen is indicated by a white arrow. The part surrounded by the broken line has been adjusted for con-trast and enlarged in the upper left of the figure. E, F. SDS-PAGE (E) and western blot (F) analyses of recombinant CD73 fused to the His-tag. Western blot analyses were performed using antibody 0614 (left) and anti–His tag antibody (right).

### 2.2. Characterization of enzymatic inhibition by chimeric antibody clone 0614-5

We converted the rat IgG2a antibody to human chimera IgG1 in order to evaluate its potential therapeutic applications (Fig. 2A). During construction of the chimeric antibody, we found that the Y78A mutation in heavy chain reduced non-specific binding in immunohistochemistry (data not shown). The impacts of mutations in this antibody will be described in detail elsewhere. We designated the human chimeric mAb 0614-5 (Fig. 2A). Next, to investigate whether this antibody had neutralizing activity, CD73 activity inhibition assay was conducted. 0614-5 did not inhibit CD73 enzymatic activity (Fig. 2B). Although this mAb does not have neutralizing activity, it could exert an antitumor effect as an armed antibody such as an antibody–drug conjugate (ADC).

**Figure 2.**
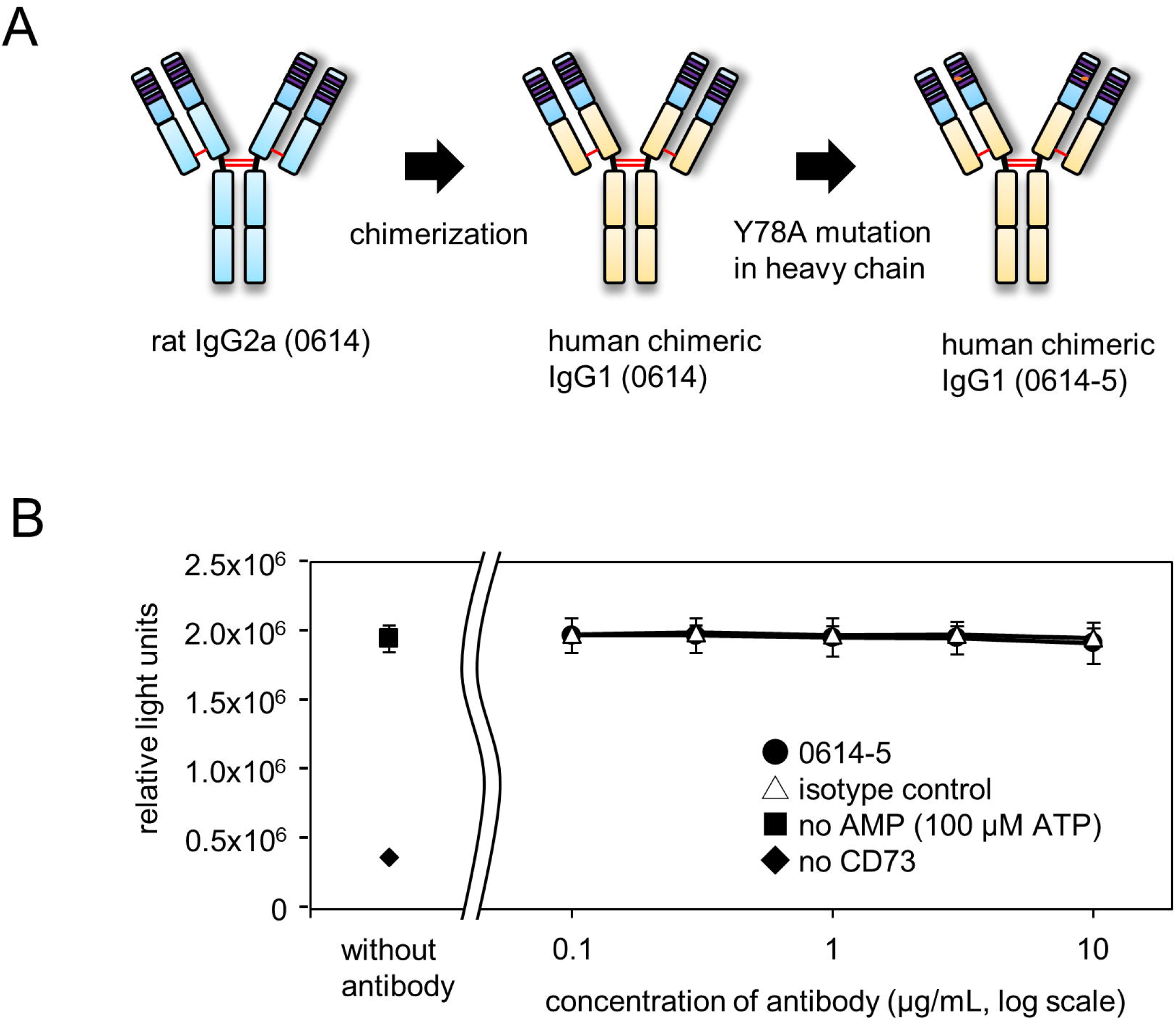
Characterization of human chimeric monoclonal antibody 0614-5. A. Schema for generating human chimeric antibody of rat 0614, 0614-5. B. Determination of inhibitory activity of mAb 0614-5. N = 3. Data are shown as means ± standard deviation.

### 2.3. Choosing cancer types for targeting by mAb 0614-5

To determine the ideal clinical application of mAb 0614-5, we screened for cancer types with high rates of CD73 positivity: either high expression in cancer cell lines or high levels in cancer tissue relative to normal samples from patients. Comparison of CD73 expression in glioblastoma, breast cancer, stomach cancer, pancreatic cancer, renal cell carcinoma, colorectal cancer, prostate cancer, and bladder cancer by flow cytometry revealed that glioblastoma (6/6), pancreatic cancer (8/8), and renal cell carcinoma (3/3) cell lines were 100% CD73-positive (Fig. 3A). Next, we compared CD73 gene expression in these cancer types using UALCAN [10], based on The Cancer Genome Atlas (TCGA) genomics data from tumor and normal samples. Among the eight cancer types, glioblastoma exhibited the highest expression of CD73 in tumor vs. normal sample (Fig. 3B). GB is among the most malignant brain tumors and is the most deadly type of cancer. Various treatments for GB are being developed, including antibody therapy [11]. CD73 is a negative prognostic factor for GB [12]. Based on these results, we decided to target GB with mAb 0614-5.

**Figure 3.**
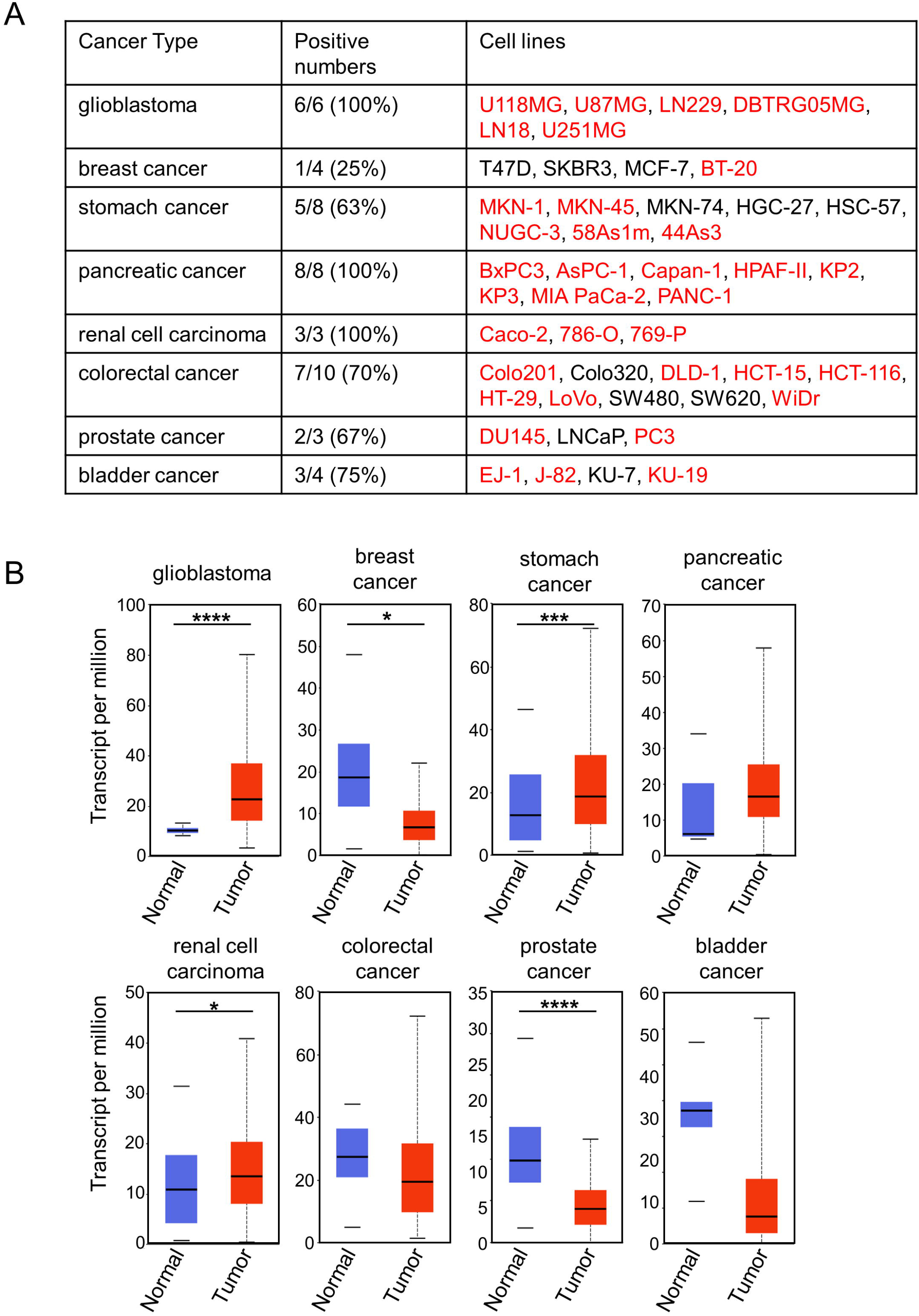
CD73 has the highest cancer specificity for glioblastoma. A. Comparison of CD73-positive rates in cell lines from different cancer types (glioblastoma, breast cancer, stomach cancer, pancreatic cancer, renal cell carcinoma, colorectal cancer, prostate cancer. and bladder cancer), as determined by flow cytometry analysis. Names of positive cell lines are shown in red. B. CD73 mRNA expression in different cancer types (glioblastoma, breast cancer, stomach cancer, pancreatic cancer, renal cell carcinoma, colorectal cancer, prostate cancer, and bladder cancer), comparing tumor (shown in red) with normal tissue (shown in blue). *P < 0.05, ***P < 0.001, ****P < 0.0001.

### 2.4. Application of clone 0614-5 antibody–drug conjugate in glioblastoma cell lines

Next, we investigated the usefulness of our 0614-5 antibody as an ADC against GB cell lines. In our in vitro cytotoxicity assays, we used six GB cell lines (U118MG, U87MG, LN229, DBTRG05MG, LN18, and U251MG). When we examined the expression of the CD73 gene in each cell line using a Cancer Cell Line Encyclopedia (CCLE) database [13], the order was as shown in Fig. 4A. We prepared mAb 0614-5 and control mAb conjugated with MMAE (0614-5-ADC and control-ADC), and then assessed in vitro cytotoxicity of control naked antibody, 0614-5 naked antibody, control-ADC, 0614-5-ADC, and free linker MMAE against GB cell lines. 0614-5-ADC exerted cytotoxicity in four cell lines (U118MG, U87MG, LN229, and DBTRG05MG) to a much greater extent than the control-ADC, and 0614-5 naked antibody had no toxic effect (Fig. 4B). In U251MG cells, which were negative for CD73 expression, 0614-5-ADC exerted no cytotoxic activity (Fig. 4A,B). Despite similar levels of CD73 gene expression, DBTRG05MG exhibited ADC-induced cytotoxicity, whereas LN18 did not (Fig. 4A,B), probably because LN18 was about 10 times less sensi-tive to MMAE than DBTRG05MG (Fig. 4C). Together, these results show that our ADC ex-erts cytotoxicity depending on the expression level of CD73. Currently, several anti-CD73 antibodies are in clinical trials (for example, AK119, BMS-986179, CPI-006, MEDI9447, NZV930, Sym024, and TJ004309) [14], and all of these antibodies block the enzymatic activity of CD73. By contrast, our antibody does not have inhibitory activity (Fig. 2B), and is thus a unique antibody that recognizes CD73 on exosomes. In the future, we hope to show that 0614-5-ADC is useful for the development of therapeutic methods targeting CD73, including combined use with checkpoint inhibitors.

**Figure 4.**
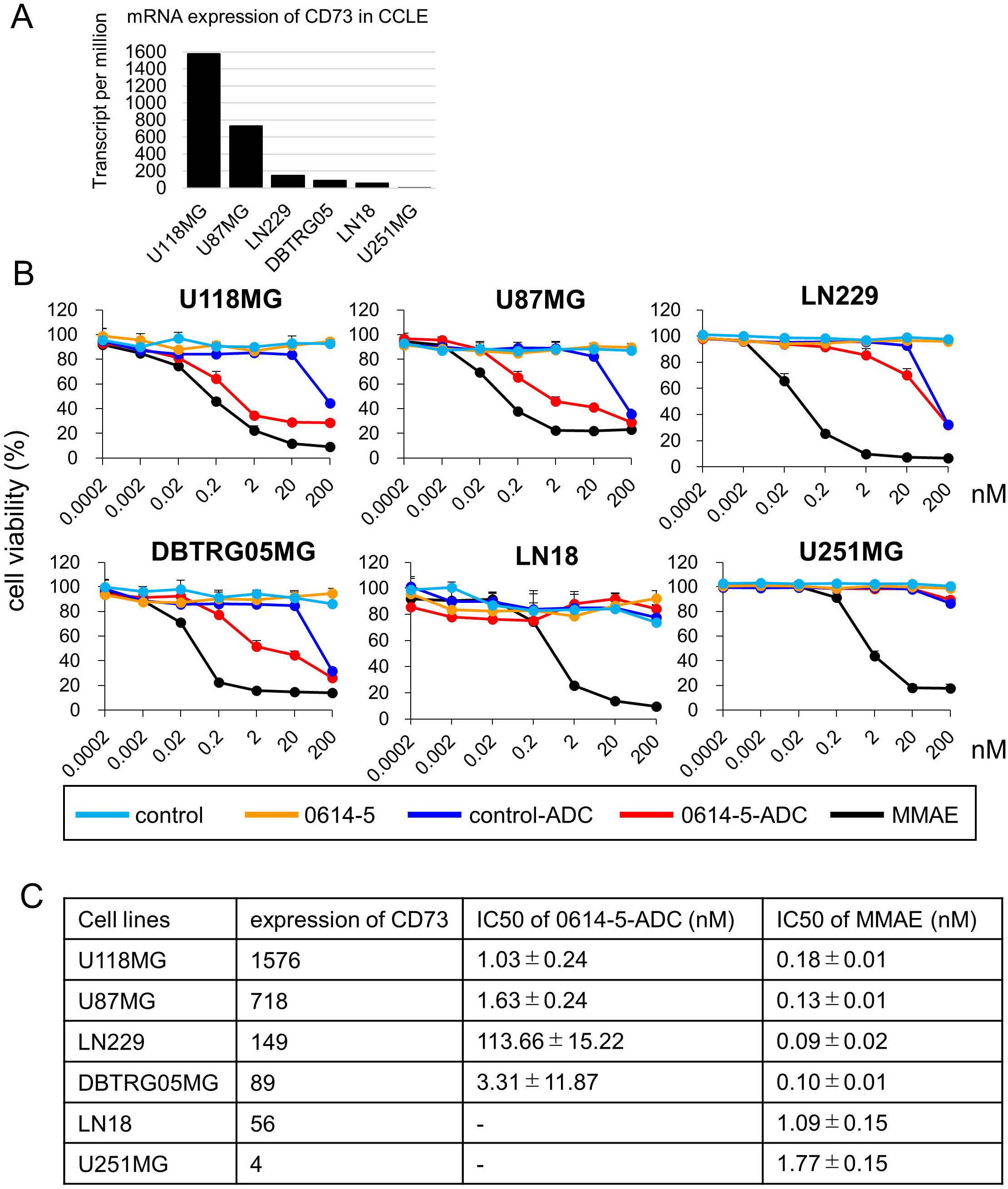
0614-5-ADC exhibits antitumor effects in GB cell lines depending on the expression level of CD73. A. mRNA expression of CD73 in GB cell lines (U118MG, U87MG, LN229, DBTRG05MG, LN18, and U251MG). B. Cytotoxicity assays performed on the GB cell lines shown in Figure 5A. N =3. Data are shown as means ± standard deviation. C. Summary of CD73 expression in Figure 5A and IC50 against GB cell lines of 0614-5-ADC or MMAE in Figure 5B.

## 3. Conclusion

In this study, we performed immunoprecipitation using magnetic beads to demonstrate that the antigen of rat antibody clone 0614, produced by exosome immunization, was CD73. Our strategy demonstrates that it is possible to identify the unknown antigens of useful antibodies. We also demonstrated that the human chimeric antibody 0614-5-ADC exerts antitumor effects on GB cell lines, dependent on the expression of CD73. Thus, 0614-5-ADC could be used to treat cancers with high levels of CD73 expression, including GB.

## 4. Materials and Methods

### 4.1. Cells and Cell culture

DLD-1, K562, U118MG, U87MG, LN229, DBTRG05MG, LN18, U251MG, and the cell lines listed in Fig. 3A were purchased from American Type Culture Collection or the Japanese Collection of Research Bioresources Cell Bank. These cell lines were cultured at 37°C under a 5% CO_2_ atmosphere in DMEM (Wako) or RPMI-1640 (Wako) supplemented with 10% FBS (Thermo Fisher Scientific) and 1% penicillin–streptomycin–amphotericin B suspension (Wako).

### 4.2. Plasmid construction

To generate chimeric antibodies, cDNAs encoding the heavy-chain variable region and kappa light-chain variable-region of antibody 0614 were PCR amplified using the following primers: heavy chain, 5’-GGATCCAACCCTTCGAATTCCACCATGGACATCAGGCTCAGC-3’ and 5’-GATGGGCCCTTGGTGCTAGCTGAGGAGACTGTGAGCATGACT-3’; light chain, 5’-GGATCGAACCCTTCGAATTCCACCATGATGGCTCCAGTTCAA-3’, and 5’-GATGGTGCAGCCACCGTACGTTTCAATTCCAGCTTGGTGCCT-3’. PCR products were cloned into vector pcDNA3.3 (Thermo Fischer Scientific) for human IgG1 expression using In-Fusion cloning technology (Clontech) as described previously[1]. For generation of 0614-5, antibody constructs were generated by inverse PCR using primers 5’-AGCACTGCCTATCCAGACTCTGTGAAG-3’ and 5’-TGGATAGGCAGTGCTACCACCAGTATT-3’ against the 0614 heavy-chain expression vector. The soluble domain (residues 27–549) of the human *CD73* gene was PCR amplified from cDNA from DLD-1 cells with an *Nde*I restriction site in the 5’ region and a *Xho*I restriction site in the 3’ region, using primers 5’-AGAAGGAGATATACATATGTGGGAGCTTACGATTTTG-3’ and 5’-GTGGTGGTGGTGGTGCTCGAGGGAAAACTTGATCCGA-3’. The PCR products were cloned into pET21a (Novagen) using InFusion cloning technology. All plasmid constructs used in this study were verified by DNA sequencing.

### 4.3. Antibodies and recombinant proteins

Rat antibody 0614 was established by exosome immunization as described previously[1]. HRP-linked 0614 rat antibody was prepared using the Peroxidase Labeling Kit-NH2 (Dojindo). Chimeric antibody expression was performed using the ExpiCHO Expression System (Thermo Fisher Scientific). Culture supernatants were purified using rProteinA Sepharose fast flow (GE Healthcare) and Superdex 200 16/600 column (GE Healthcare).

Purified monoclonal antibodies were buffer-exchanged into PBS, concentrated using Vivaspin Turbo 15 30-kDa centrifugal filter units (Sartorius), and stored at 4°C until use. Recombinant CD73 protein was overexpressed in *E. coli* Rosetta (DE3) cells. The insoluble fraction was solubilized and purified by Ni-NTA agarose resin (Invitrogen) under denaturing conditions (8 M urea). Purified denatured CD73 was refolded by the rapid dilution method using a refolding buffer consisting of 20 mM Tris–HCl pH 8.0, 0.5 M L-arginine, and 10% glycerol. Renatured CD73 protein was purified by Superdex 75 16/600 column. Purified CD73 was buffer-exchanged into PBS, concentrated using Amicon ultra 4 10-kDa centrifugal filter units (Millipore), and stored at 4°C until use. Purity of proteins was determined by SDS-PAGE using 4–15% Mini-PROTEAN TGX gel (Bio-Rad). The C-terminal purification His-tag was not removed.

### 4.4. Isolation of antigen of antibody 0614 from exosome sample

0614 antibodies were biotinylated with EZ-Link NHS-LC-Biotin (Thermo Fisher Scientific). Biotinylated antibody 0614 (10 μg) and 0.1 mg of streptavidin-conjugated magnetic beads (FG beads, Tamagawa Seiki) were reacted at 4°C for 1 hour, and then washed by magnetic separation. Exosomes from CRC lines or exosomes from blood cell lines (as a negative control) were reacted for 2 hours with 0.1 mg of antibody 0614-conjugated magnetic beads in PBS-T. To crosslink antibodies and antigens, 5 nmol of DTSSP was reacted at room temperature for 30 minutes, and then the reaction was quenched with 50 nmol of Tris for 15 minutes. Sample collected by magnetic beads was solubilized by RIPA buffer (Wako) and then analyzed by SDS-PAGE and western blotting. SDS-PAGE was performed using 7.5% Mini-PROTEAN TGX gel (Bio-Rad), and the gels were stained with Oriole fluorescent gel stain (Bio-Rad). For western blotting, samples were transferred to PVDF membranes (Merck), and the membranes were blocked for 15 minutes with StartingBlock (TBS) Blocking Buffer (Thermo Fischer Scientific). The blocked membranes were washed three times TBS with 0.1% Tween 20 (TBS-T) and then incubated with HRP-linked antibody 0614 (2 μg/mL) at room temperature for 1 hour. After the membranes were washed five times with TBS-T, the proteins were visualized using Chemi-Lumi One Super (Nacalai Tesque).

### 4.5. Identification of the antigen of antibody 0614

For 2D-PAGE, acetone precipitation was performed, and 20-μL samples were mixed with 80 μL ice-cold acetone and stored at −20°C for 3 hours. The samples were then centrifuged at 20,000 *g* for 10 minutes at 4°C, and after the supernatant was discarded, the sediment was dried. The samples were resuspended with 15 μL 1% n-octyl-β-D-glucoside / 8 M urea / 2 M Thiourea and centrifuged at 20,000 *g* for 5 minutes at 4°C. Supernatants (10 μl) were subjected to 2D-PAGE by Auto2D (SHARP). Gel staining was performed using the Silver Stain MS Kit (Wako). Shotgun proteome analysis was performed by Medical ProteoScope. Target protein spots and control spots were excised from the gel and enzymatically digested in-gel as previously described [15]. Gel pieces were destained using the Silver Stain MS Kit (Wako) and washed with acetonitrile. Samples were alkylated with DTT and iodoacetamide, and then trypsinized for 16 hours at 37°C. Trypsin-digested peptides were extracted with 25 mM ammonium hydrogen carbonate, 5% formic acid, and acetonitrile and then dried. Liquid chromatography was performed on a fully automated Ultimate U3000 Nano LC System (Thermo Fisher Scientific). MS/MS analysis was performed on a Q Exactive Orbitrap (Thermo Fisher Scientific). Data were analyzed using Proteome Discoverer (ver. 2.2) (https://www.thermofisher.com/jp/ja/home.html), Mascot (Matrix Science), and an in-house database.

### 4.6. Confirmation of the antigen of antibody 0614 as CD73

The antigen of antibody 0614 was confirmed by western blotting. Recombinant CD73 with a C-terminal His-tag was loaded onto SDS-PAGE gel, separated, and then transferred to PVDF membranes (Bio-Rad). Membranes were blocked for 5 minutes with Bullet Blocking One for western blotting (Nacalai Tesque). The membranes were washed three times with TBS-T. The blots were then incubated with antibody 0614 (10 μg/mL) or anti-His tag antibody (1:10000, ProteinTech) at room temperature for 1 hour. The membranes were washed three times with TBS-T and then incubated with HRP-conjugated anti–rat IgG (1:5000; Jackson Immuno Research) or anti–mouse IgG (1:5000; CST) at room temperature for 1 hour. Can Get Signal solution (Toyobo) was used to reduce background noise. After the membranes were washed three times with TBS-T, proteins were visualized using ECL prime (GE Healthcare).

### 4.7. CD73 activity inhibition assay

The CD73 activity inhibition assay was conducted with reference to a previous study [16]. The indicated concentrations of antibody 0614-5 or isotype control antibody were incubated with refolded recombinant CD73 either prior to or after substrate (adenosine monophosphate [AMP]) addition. Adenosine triphosphate (ATP) levels were measured by CellTiter-Glo 2.0 (Promega). An AMP-free sample containing 100 μM ATP was used as a positive control for ATP measurement, and a CD73-free sample was tested as a negative control.

### 4.8. Flow cytometry

Flow cytometry was conducted as previously described [1]. Cultured cells were stained with mAb 0614 or human chimera MAb 0614-5 as the primary antibody and Alexa Fluor 488– or 647–conjugated anti-rat or human-Ig polyclonal antibody (Thermo Fisher Scientific) as the secondary antibody. Stained cells were analyzed on a Guava easyCyte 10HT (Merck Millipore). Dead cells were stained using propidium iodide (PI) (Thermo Fisher Scientific). Data were analyzed using the FlowJo software (Tree Star).

### 4.9. Database analysis

CD73 gene expression in normal and tumor samples from the TCGA cancer genomics data was analyzed using UALCAN[10]. Overall survival and disease-free survival in the high NT5E expression and low-expression groups from the TCGA data were analyzed using GEPIA2[17].

### 4.10. *In vitro* cytotoxicity assay

The drug payload MMAE was purchased from MedChem Express. Generation of ADC was conducted as described previously [18]. Cells were plated at 3,000 cells/well in 96-well plates and cultured for overnight. After media was replaced, the indicated con-centrations of naked antibodies, ADCs, or free MMAE were added, and the sample was incubated for 72 hours. Cell viabilities were monitored using the Cell Counting Kit-8 (Dojindo).

## Author Contributions

Conceptualization: T.A., S.S., Y.M. and M.Y.; Methodology: T.A., S.S., and M.H.; Analysis, Investigation, and Data Curation: T.A., S.S., H.T., M.H., and S.H.; Writing—Original Draft Preparation: T.A.; Writing—Review and Editing: T.A., and M.Y.; Supervision: Y.M. and M.Y.; Funding Acquisition: T.A., Y.M., and M.Y. All authors have read and agreed to the final version of the manuscript.

## Institutional Review Board Statement

Not applicable.

## Informed Consent Statement

Not applicable.

## Data Availability Statement

The raw data presented in this study are available on request from the corresponding author.

## Acknowledgements

The authors thank Matsumura and Yasunaga laboratory members for helpful discussion, and M. Shimada for their secretarial support.

---

**Figure S1. Screening of hybridoma cells containing antibodies by flow cytometry**

Flow cytometric analysis of colorectal cancer cell line DLD-1 and myeloma cell line K562 by culture supernatant of hybridoma producing antibody 0614.

