## Supplementary figures and images for "Identification of CD73 as the antigen of an antigen-unknown monoclonal antibody established by exosome immunization, and its antibody-drug conjugate exerts an antitumor effect on glioblastoma cell lines"

### FigureS1

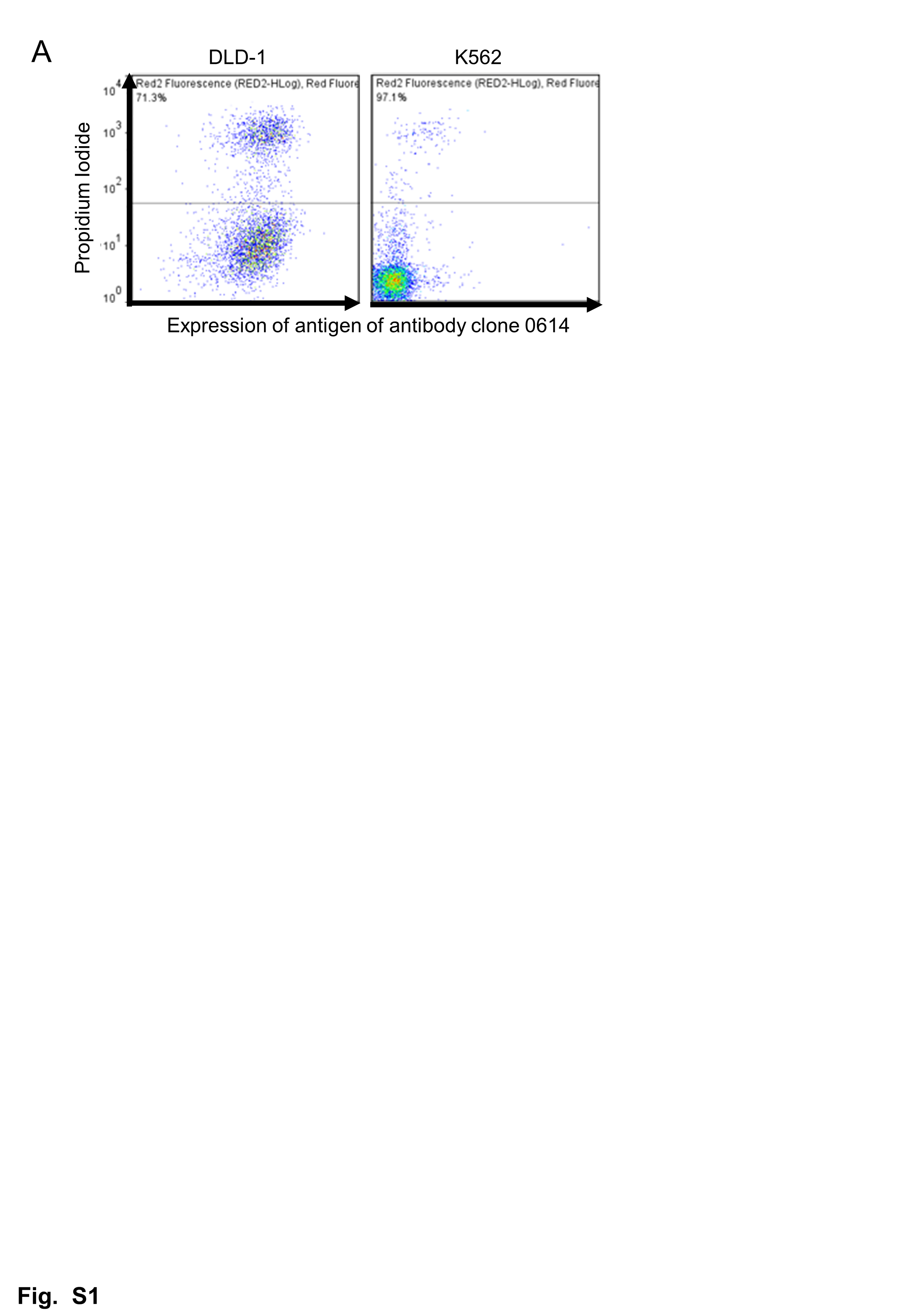
